# Mechanical Ventilation induced DNA Damage and P21 in an Acute Aging Model of Lung Injury

**DOI:** 10.1101/2022.03.08.483505

**Authors:** Franck J. Kamga Gninzeko, Cindy K. Tho, Michael S. Valentine, Emily N. Wandling, Rebecca L. Heise

## Abstract

Acute respiratory distress syndrome (ARDS) is a form of acute lung injury which leads to a paucity of oxygen. To remedy ARDS, patients are put on mechanical ventilators; however, the stretch resulting from the mechanical ventilator can lead to ventilator-induced lung injury or VILI. VILI is exacerbated by age. The combined effects of ARDS and mechanical ventilation can result in a hostile environment which may lead to senescence (stable cell cycle arrest). The role of senescence in VILI is poorly understood. Senescence is characterized by increased cyclin-dependent kinase inhibitors P16 and P21; however, P21 has been shown to occur in early senescence. We hypothesized that mechanical ventilation would lead to DNA damage and senescence-like phenotype. Both in vivo and in vitro models of VILI were used to investigate senescence and its mechanism in VILI. Mechanical ventilation increased senescence-associated markers such as DNA damage marker characterized by ɣH2AX, P21, senescence-associated secretory phenotype IL6, and decreased proliferation. Moreover, mechanical ventilation led to increased apoptosis. Lung sections were stained for KRT8 proteins, markers of transiently differentiated alveolar type 2 (AT2) cells which were reported to be more prone to DNA damage. Age and mechanical ventilation increased KRT8 positive cells. Finally, we probed a potential mediator of the stretch induced senescence *in vitro* by inhibiting P38-MAPK, which can be activated by DNA damage response, leading to increased P21. Inhibiting P38 decreased in P21 but did not decrease ɣH2AXThese findings suggest that mechanical ventilation may lead to a senescence-like phenotype involving the P38-MAPK pathway.

## Introduction

Acute respiratory distress syndrome (ARDS) is characterized by a leaky alveolar barrier and inflammation. ARDS has a high mortality rate and disproportionally affects the elderly population with a higher mortality and incidence rate [1]. There are signs of hindered gas exchange amongst ARDS patients [2], leading to mechanical ventilation. Despite the benefits of mechanical ventilation at mitigating the effects of ARDS and facilitating the gas exchange, the constant strain and stress created by the former due to the air moving in and out of the alveoli may worsen the injury and lead to ventilation-induced lung injury, VILI [3]. VILI is characterized by biotrauma due to the increased inflammation; volutrauma due to the overextension of the alveoli caused by air going in and out of the lungs during ventilation; and atelectrauma caused by the constant collapse and reopening of the alveoli [4]. These characteristics of VILI, especially biotrauma and volutrauma, can result in cascades of events that can cause the alveoli cells to become damaged and change phenotype, creating a hostile environment. The hostile environment created by VILI coupled with inflammation observed in ARDS may lead to senescence.

Senescence is an irreversible terminal cell cycle arrest process in which proliferating cells stop responding to replication-promoting stimuli. However, evidence suggests that senescence can occur prematurely in response to injury [5]. Because of its heterogenicity across different tissues and conditions, senescence is a complex phenomenon that is poorly understood. Furthermore, more is known about senescence in chronic lung diseases than acute. In senescence, cyclin-dependent kinase inhibitors such as P21 and P16 are upregulated. While P21 plays an essential role in early senescence, P16 is accumulated at a later stage of senescence [6]. There is also evidence that P21, not P16, is upregulated with VILI [7]. Senescent cells also produce autocrine and paracrine signaling collectively known as senescence-associated secretory phenotype, SASP. SASP mediators include pro-inflammatory cytokines such as IL-6, which can exacerbate inflammation [8].

Though some parts of the senescence mechanism have been investigated, more still needs to be done to fully understand the mechanism of senescence in VILI and other diseases. One of the mechanisms of senescence is the DNA damage response pathway which could be activated via mechanotransduction. Cyclic stretch like the one generated by mechanical ventilation can break DNA strands down via MAPK activation [9], leading to upregulation of DNA damage proteins such as ɣH2AX [10]. Activation of ɣH2AX, in turn, will lead to the activation of P21 [11]. P38-MAPK is also known to activate the P53-P21 pathway resulting in cellular senescence [11]. However, this pathway has not been investigated in the context of senescence in acute diseases or VILI.

In this research, we are investigating whether mechanical ventilation promotes senescence-like phenotypes. We also examine whether stretch induces DNA damage response, leading to senescence. Finally, we are using a P38-MAPK inhibitor to understand the mechanism of senescence in VILI. We hypothesized that mechanical ventilation would lead to DNA damage and senescence-like phenotype via P38-MAPK activation, and blocking it will reduce this phenotype.

## Methods

### Animal

Male young (8-10 weeks) and old (20-22 months) C57BL/6 mice were acquired from the National Institute on aging and housed at the VCU vivarium. All procedures performed on the mice were approved by VCU Institutional Animal Care and Use Committee (IACUC). Mice were mechanically ventilated at 0 PEEP and 35 and 45 cmH2O for young and old mice, respectively, for 2 hours. Lung mechanics were measured at 30-minute increments for the duration of ventilation.

### Sample Collection

At the end of the 2-hour mechanical ventilation, blood was collected from the vena cava and centrifuged to obtain plasma. As previously described, gravity-assisted bronchoalveolar lavage (BAL) was performed [12]. Briefly, PBS was instilled into the mice lungs and collected and repeated twice. The BAL fluid (BALF) was then centrifuged, the supernatant was transferred onto a new tube for later processing. The cells were resuspended then cytospun onto a microscope slide. WBC analysis was performed by counting 300 cells while refraining from the peripheral edges of the microscopy region. The different immune cells were distinguished and quantified.

### In vitro cell stretch

Human small airway epithelial cells (SAEC) were purchased from Promocell (C-12642) and cultured according to their recommendations with a supplied SAEC growth media. SAEC were plated on Bioflex culture plates (Flexcell international Corp., BF-3001) for 48 hours for acclimation. Then, media change was performed; some wells received DMSO vehicle and others SB203580 (a P38 inhibitor) for 30 minutes. Media change was performed again to rid the wells of the vehicle and inhibitor. The cells were then stretched at 20% change in surface area for 24 hours and at 0.33 Hz to cause injury.

### qPCR

For lung tissues, small lung fragments of about 50 mg were homogenized with Trizol (ThermoFisher, 15596026). RNA extraction was performed as recommended by the manufacturer. Briefly, after homogenization, 200 ul of chloroform was added per 1 ml of Trizol. Phase separation was performed via centrifugation, then the clear phase, which contained RNA, was then separated. *In vitro* experiments, cells were lysed using RLT buffer to obtain a cell lysate containing RNA. RNA isolation of both tissues extracted RNA, and cell lysate was performed using the Qiagen RNeasy kit. cDNA conversion was performed using a Bio-Rad iScript cDNA synthesis kit (Bio-Rad, 1708891). cDNA and Bio-Rad SsoAdvanced universal SYBR green (Bio-Rad, 1725274) was used for gene amplification. Primers: human 18s F: 5’-TAACCCGTTGAACCCCATTC-3’ R: 5’-TCCAATCGGTAGTAGCGACG-3’, human P21 F: 5’-TGTCCGTCAGAACCCATGC-3’ R: 5’-AAAGTCGAAGTTCCATCGCTC3’, mouse 18s: F: 5’-GCAATTATTCCCCATGAACG-3’ R: 5’-GGCCTCACTAAACCATCCAA-3’, mouse P21 F: 5’-GACAAGAGGCCCAGTACTTC-3’ R: 5’-GCTTGGAGTGATAGAAATCTGTC-3’

### Inflammatory Cytokine Analysis

IL6 ELISA (R&D system, DY 406) was performed according to the manufacturer’s protocol. Briefly, 96-well ELISA plates were coated using a captured antibody overnight. The next day, the plate was washed (0.05% Tween 20 in PBS) and blocked (R&D system, DY995) at room temperature using the reagent diluent for 2 hours. Standards and samples were loaded into the plates and incubated at room temperature for 2 hours. The plates were then washed, and the detection antibody was added for 2 more hours. The plates were then incubated with Streptavidin-HRP and substrate solution solutions, respectively, for 20 minutes, each with washes in between. Diluted sulfuric acid at the concentration recommended by the manufacturer was used to stop the reaction. The plates were read at 450 nm with wavelength correction.

### Immunofluorescent staining

Immunofluorescent staining was performed according to the antibody manufacturer protocol with modifications. Briefly, when stretch experiments concluded, the cells were fixed then permeabilized. The cells were blocked for 2 hours at room temperature using 10% normal goat serum or BSA when appropriate. The slides were then incubated overnight at 4C in primary antibody. The next day, the slides were rinsed then incubated with a secondary fluorophore antibody for an hour. For tissue embedded sections, the slides were deparaffinized, and antigen retrieval was performed before the permeabilization. Primary antibodies: mouse anti-gamma H2AX (Novus Biologicals, NB100-74435), rabbit anti-Ki67/MKi67 (Novus Biologicals, NB500-170SS), rabbit anti-SP-C (Bioss, bs-10067R), rat anti-KRT8 (CreativeBiolabs, CBDH1469). Goat anti-Rat IgG (H+L) Alexa Fluor 555 (ThermoFisher Scientific, A-21434), Goat anti-Rabbit IgG (H+L) Alexa Fluor 647 (ThermoFisher Scientific, A-21245), Goat anti-Mouse IgG (H+L) Alexa Fluor 488 (ThermoFisher Scientific, A-28175).

### Tissue Protein extraction

cytoplasmic and nuclear proteins were extracted according to the Millipore sigma protocol with minor modifications. Briefly, tissues were lysed using hypotonic buffer (10 mM HEPES, pH 7.9, with 1.5 mM MgCl2 and 10 mM KCl) containing Dithiothreitol (DTT), protease and phosphatase inhibitors. Then the tissues were homogenized and centrifuged at 10,000 g for 20 minutes. The supernatant, which contained cytoplasmic proteins, was preserved. The crude nuclei pellet was resuspended in 140 ul of extraction buffer containing DTT, protease, and phosphatase inhibitors. The solution was gently shaken for 30 minutes then centrifuged for 5 minutes at 20,000 g. The supernatant was transferred to a new tube and chilled at −70 C for later processing.

### Western Blot

Western blot was performed according to the manufacturer’s protocol. Briefly, BCA was performed on the freshly isolated proteins using a Pierce BCA protein assay kit (Sigma, 23227) to determine the protein concentration. Electrophoresis was performed using 30 ug of proteins loaded into each 10-well gel; proteins were then transferred from the gel onto a membrane. The membrane was blocked for 1 hour, followed by primary antibody incubation overnight at 4 C. The next day, the membrane was rinsed and incubated in a solution of HRP bound secondary antibody for 1 hour, followed by rinsing and incubation in Pierce ECL western blotting substrate (ThermoFisher, 32106) for 5 minutes and imaged. Bands densities were analyzed using BioRad Image Lab software. Rabbit anti-histone H3 (Novus Biologicals, NB500-171), mouse anti-gamma H2AX (Novus Biologicals, NB100-74435), anti-rabbit IgG HRP-linked antibody (Cell Signaling Technology, 7074S), anti-mouse IgG HRP-linked antibody (Cell Signaling Technology, 7076S).

### TUNEL staining

Click-iT™ Plus TUNEL assay for in situ apoptosis detection with Alexa Fuor™ 647 was performed according to the manufacturer protocol (ThermoFisher Scientific, C10619). Briefly, slides were deparaffinized using xylenes, decreasing percentages of ethanol, saline solution, and PBS. The tissue slides were fixed and permeabilized using 4% paraformaldehyde and proteinase K. Then, TdT reaction was performed by incorporating EdUTP into dsDNA strand breaks. Finally, fluorescence Click-iT™ plus reaction was performed to detect EdUTP; then the slides were mounted using ProLong™ Gold Antifade Mountant with DAPI (ThermoFisher Scientific, P36931).

### Statistical analysis

Subjects were randomly assigned to different groups. Power analysis was estimated based on previous work. A minimum of 3 mice per group was used. One-way and two-way ANOVA followed by posthoc Tukey’s multiple comparison test were performed when appropriate to compare multiple groups. A T-test was also performed to compare two groups. GraphPad Prism 6 was used for statistical analysis.

## Results

### Mechanical Ventilation and Age Cause Structural Damage and Increased Polymorphonuclear Cells in BALF

To assess the effects of mechanical ventilation on the lung structure and physiology, lung histology, lung mechanics perturbation maneuvers, and PMN (Polymorphonuclear leukocytes) count were performed. H&E-stained lung sections showed that high-pressure MV caused increased structural damage on the lungs of both young and old mice. Furthermore, when considering age alone, old mice inherently showed more significant structural damage signs than young mice without mechanical ventilation (figure 1A). BCA results showed increased protein in the BALF with age and ventilation. These results suggest that mechanical ventilation and age are cofactors for lung injury (figure 1B).

**Figure 1:**
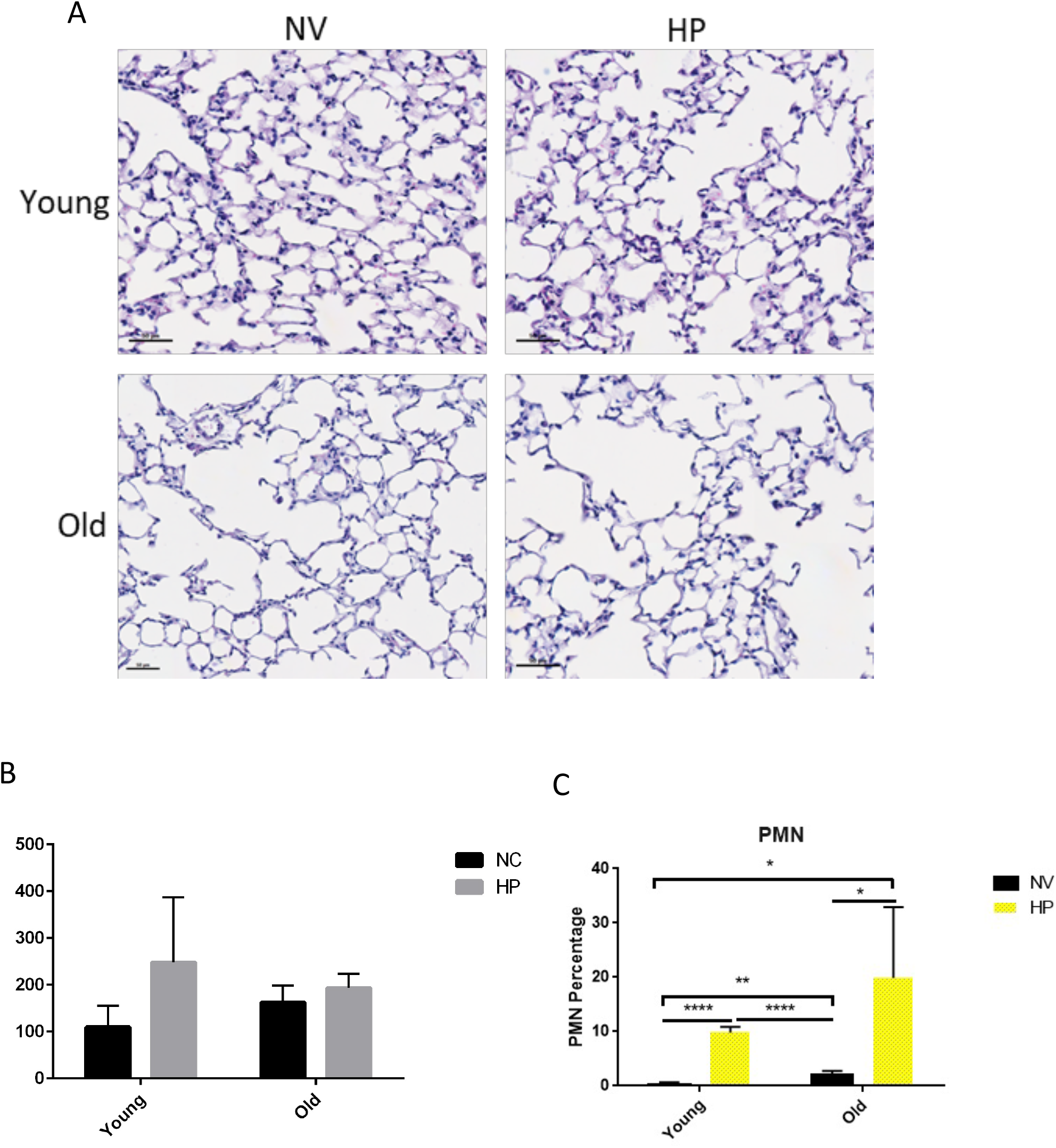
A: H&E staining of lung histology showing increased structural damage with age and mechanical ventilation B: BCA (N=3-5) increase protein in BALF with age and mechanical ventilation though not significant C: PMN (N=4-5) of mechanically ventilated mice showing increased PMN recruitment with mechanical ventilation and age. Scale bar 100 um. NV: non-ventilated and HP: high pressure. * p < 0.05, ** p < 0.01 and **** p < 0.0001.

Similarly, the count of PMN on BALF showed a significant increase in PMN intrusion in the alveoli space with mechanical ventilation in both age groups. At baseline, without mechanical ventilation, old mice showed significantly higher number of PMN in the BALF compared to the young non-ventilated. However, in the old group who did not receive mechanical ventilation, we did not see much increase in PMN compared to the young non mechanically ventilated (figure 1C).

#### Lung Mechanics

From the PV-loops, we first noticed in both young and old mice that the loops have shifted up and a bit to the left after 30 minutes of ventilations, suggesting an increase in compliance. After one hour on the ventilator, both age group PV-loops were shifted back down and to the right, closer to the baseline’s loop. However, two hours after mechanical ventilation, in the young mice group, there was a shift down and to the left compared to baseline compliance suggesting a decrease in compliance. After that initial increase in compliance, the loop returned to baseline in the old subjects (figure 2).

**Figure 2:**
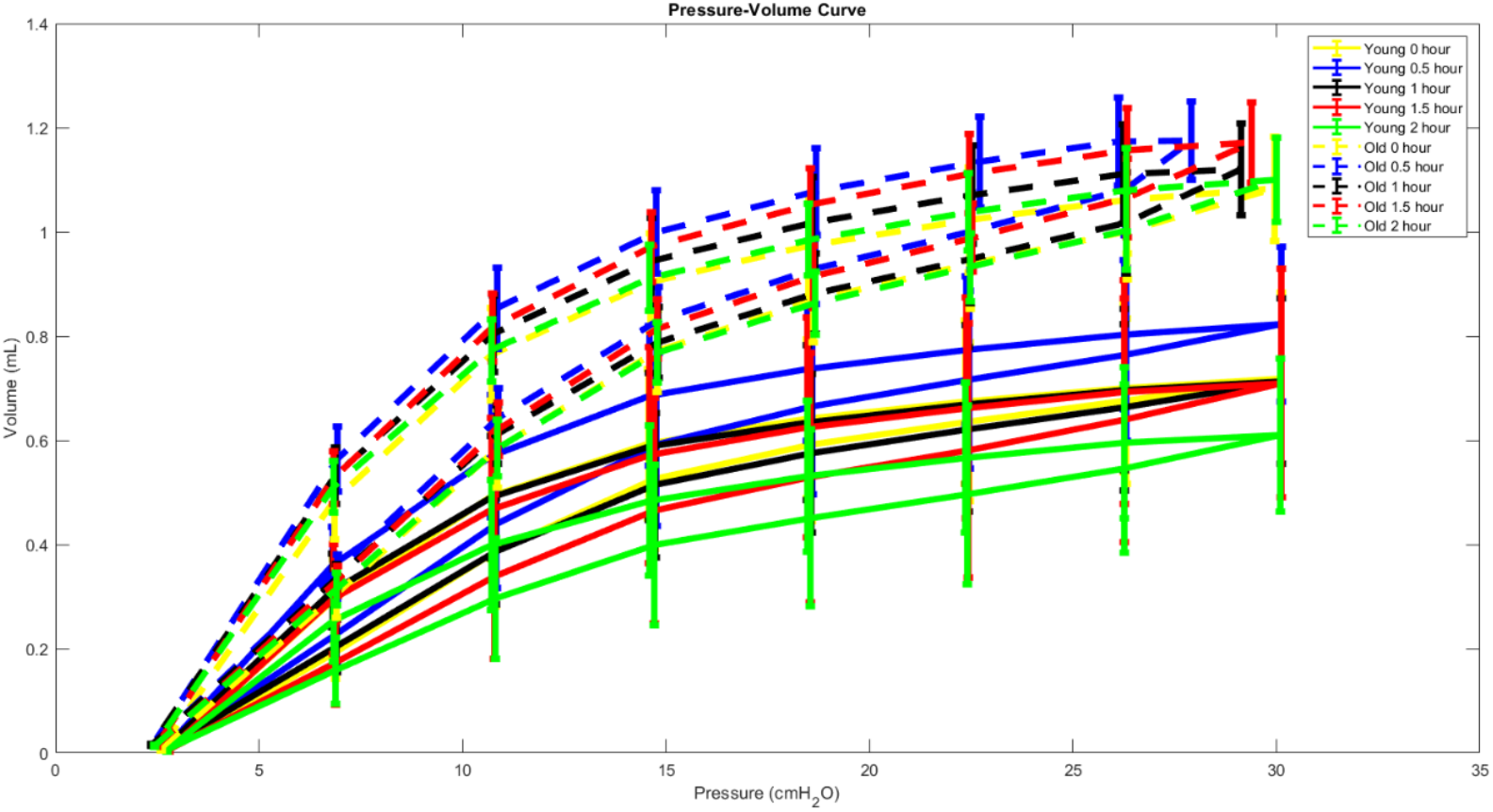
Single compartment static lung compliance showing changes in the lung mechanics of mechanically ventilated mice with age. N = 3-5 mice

##### TUNEL

TUNEL was used to detect apoptosis in the fixed lung tissue postmortem. There was an increase in TUNEL positive cells with mechanical ventilation in the young mice group. Moreover, non-ventilated old mice showed signs of apoptosis though not in a considerable amount. However, old mice showed more apoptotic cells than other groups when mechanically ventilated. These results suggest that the high-pressure mechanical ventilation settings used in this study caused damage at the tissue level, which changed the lungs’ mechanics, and damage at the cellular level, which manifested by increased apoptosis in both you and old groups (figure 3).

**Figure 3:**
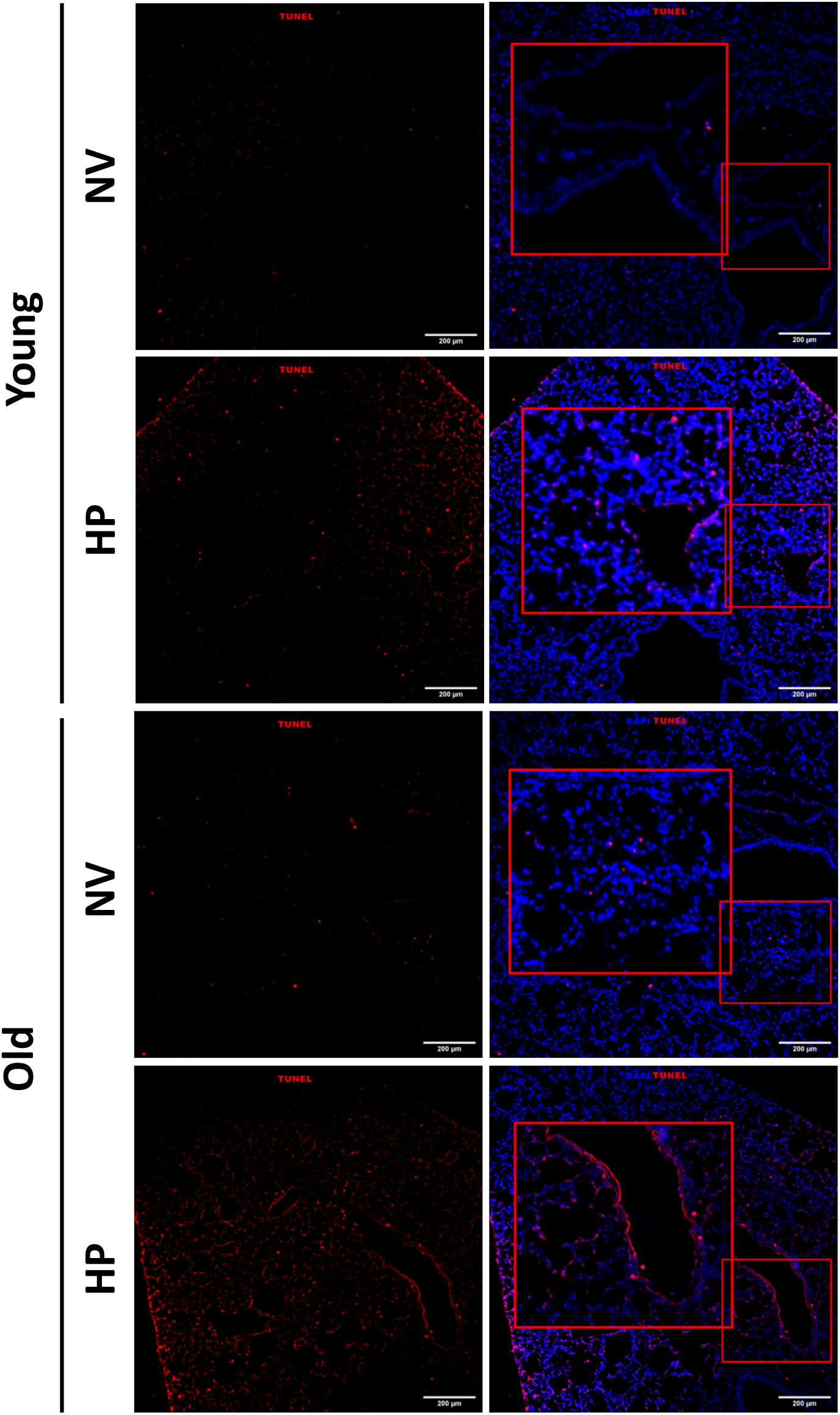
TUNEL staining showing increased apoptosis in the lungs of young and old mice with mechanical ventilation. Young mice also showed signs of apoptosis without mechanical ventilation. NV: non-ventilated and HP: high pressure.

### Presence of Senescence-like phenotypes in Mechanical Ventilation induced Acute Lung Injury

#### DNA Damage increases with age, and HP

One known pathway leading to cellular senescence is the DNA damage response pathway. In this experiment, after mechanical ventilation, we probed for ɣH2AX, a DNA damage marker in the homogenized bulk lung nuclear protein. Mechanical ventilation of both young and old mice yielded an increase of ɣH2AX, a sign that the mechanical force generated by the mechanical ventilator is transduced from the extracellular space in the cytoplasm and the nucleus of the cells causing damage to the nucleus. Moreover, in the old mice population, there was an increase in ɣH2AX with no mechanical ventilation compared to the young mice who were not mechanically ventilated, suggesting that old mice inherently show signs of DNA damage probably due to age (figure 4 A).

**Figure 4:**
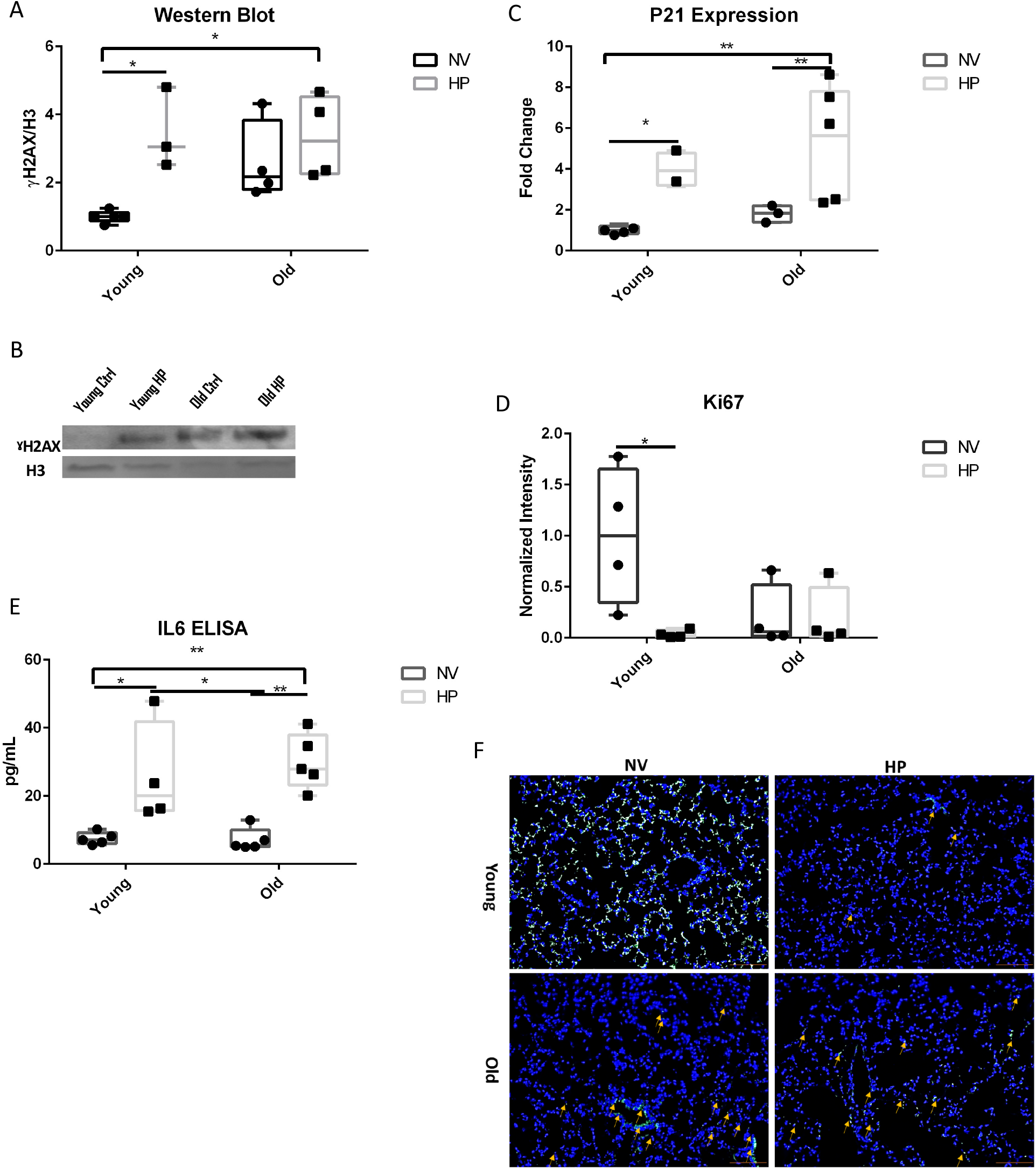
A: ɣH2AX increased with ventilation in both age groups, however not with age alone. B: Representative western blot of ɣH2AX C: P21 increased with ventilation as well. D: Decreased Ki67 with age and with mechanical ventilation. E: BALF IL6 increased as a result of mechanical ventilation. F: Representative images of Ki67 staining. Scale bar 100 um. NV: non-ventilated and HP: high pressure. N = 4-6. * p < 0.05, ** p < 0.01.

#### HP is correlated with P21 gene expression, SASP, and decreased proliferation

P21, a protein involved in the cell cycle, was analyzed. There was an increase in P21 gene expression with mechanical ventilation. Young and old mice that were mechanically ventilated had a significant increase in P21 compared to the young nonventilated. Mechanically ventilated old mice also showed a significant increase in P21 compared to the old non-ventilated. These results suggest that mechanical ventilation plays a role in P21 regulation. However, we did not observe a significant increase in P21 gene expression with age alone; this could be because P21 is an early marker of senescence (figure 4B).

Ki67, a proliferation marker, did not increase with age or mechanical ventilation. Ki67 level was significantly lower in the young mice when mechanically ventilated compared to the young non-ventilated. This is consistent with the P21 marker increase suggesting that mechanical ventilation and age initially lead to increased P21 gene expression and the lack of proliferation (figure 4C).

IL6, a pro-inflammatory marker part of the SASP, was also probed in the BALF. It has been previously shown that when cells become senescent, they produce cytokines which include IL6 [8]. Mechanically ventilated mice showed an increase in IL6 level (figure 4D); this is on par with P21 gene expression groups. In all, these results suggest that increased DNA damage, high P21 gene expression levels, and reduced Ki67 (proliferation), all hallmarks of cellular senescence, also lead to increase SASP.

### Age and mechanical Ventilation lead to an Increase in Cells expressing keratin 8 (KRT8+)

There are mainly two cell types in the alveolar epithelium, alveolar type 1 and type 2 (AT1 and AT2). It has been shown that due to mechanical cues and in certain diseases, AT2 differentiate into AT1[13]. It has also been shown that before differentiating to AT1 due to various stimuli, AT2 cells go through a transient state which is positive for Krt8 [14]. These transient KRT8+ cells are also more prone to DNA damage [15]. To test if mechanical ventilation resulted in increased KRT8+ cells, we stained the lungs of the mice with anti-KRT8 antibodies. Compared to the young non-ventilated mice, there was an increase in KRT8+ cells in the young mechanically ventilated mice. Moreover, there was an increase in KRT8+ cells in the old mice group, regardless of ventilation status, compared to the young non-ventilated mice. The increase in KRT8+ cells (which are more prone to DNA damage) with age and mechanical ventilation could explain why there was an increase in DNA damage in those groups compared to the non-ventilated young mice (figure 5).

**Figure 5:**
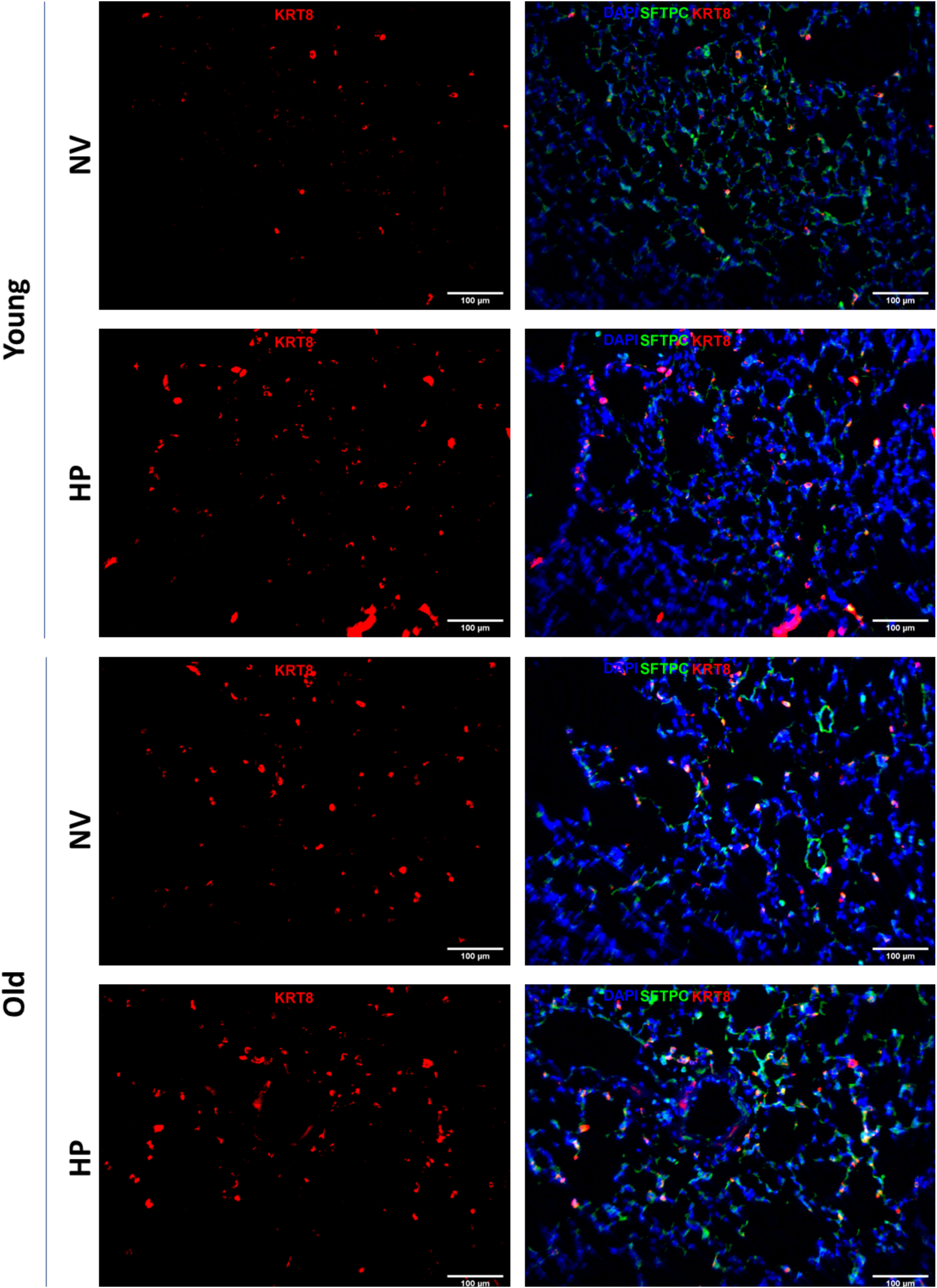
Immunofluorescence staining of lung section for both young and old mice. Staining for KRT8 (red), a marker of transiently differentiated AT2, and SFTPC (green), an AT2 marker. DAPI (blue) was used as a counterstain. The staining reveals an increase in KRT8 positive cells with age and mechanical ventilation and a decrease in SFTPC in all groups compared to young non-ventilated. NV: non-ventilated and HP: high pressure.

### Cell stretch-induced senescence markers in human SAEC

We first wanted to know if human SAEC undergoes DNA damage in response to cyclic stretch, potentially leading to cellular senescence. Compared to the static group, there was more DNA damage manifested by an increase ɣH2AX in the stretch SAEC group (figure 6 A). Similarly, there was an increase in P21 in stretch SAEC compared to the static group (figure 6B). This is consistent with the results obtained in the mechanically ventilated old mice; The SAEC were from patients aged 51-66 years. As in old mice, there was an increase in ɣH2AX with mechanical stretch compared to the static group. In addition, it was only when mechanically ventilated that the old mice showed an increase in P21.

**Figure 6:**
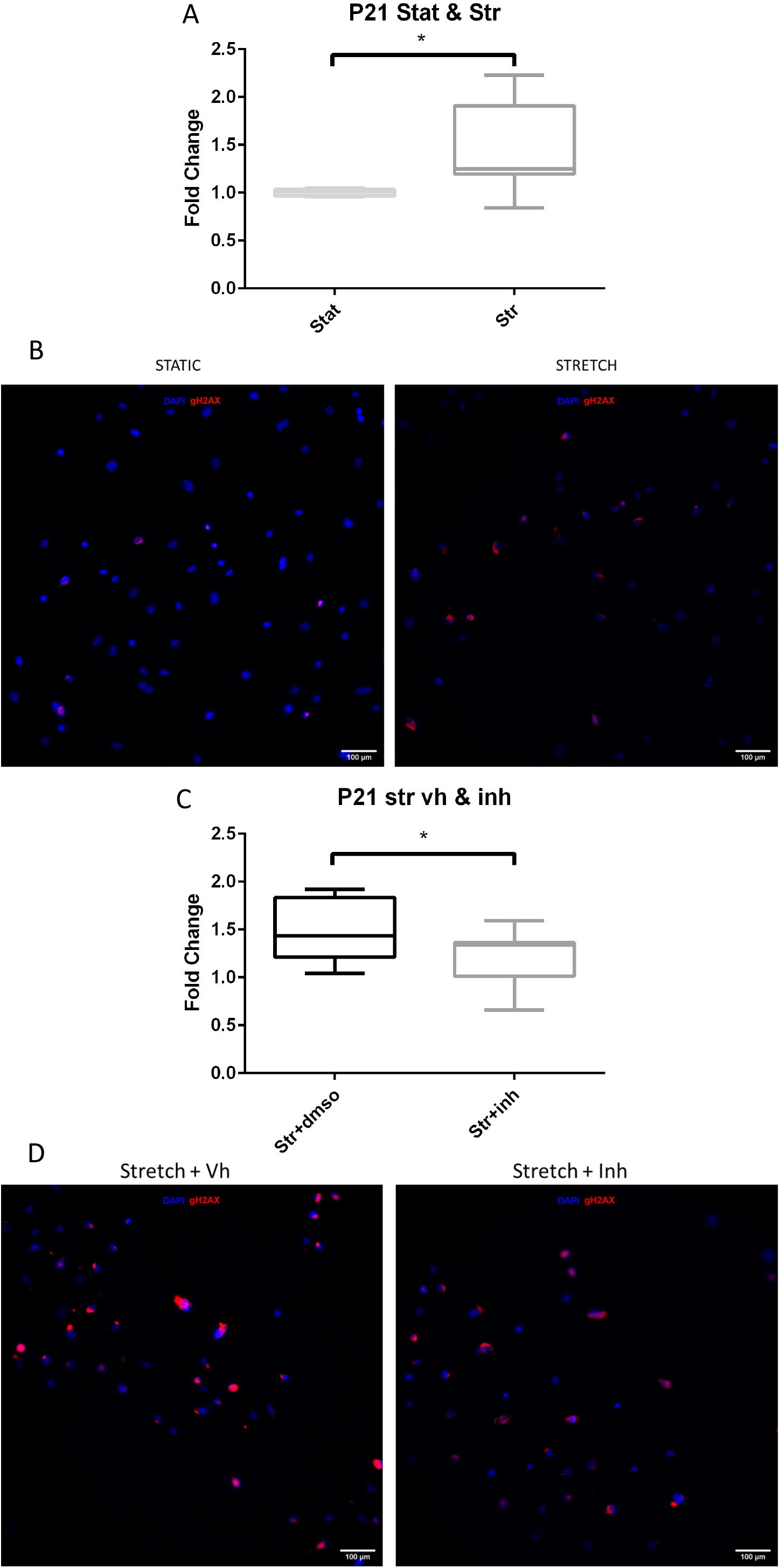
Gene expression and immunofluorescent staining of human SAEC cells at static and stretch conditions and with DMSO and P38 inhibitor A&B: Stretch Caused Increase P21 and DNA Damage in SAEC, C&D: P38 inhibition reduces P21 but does not affect DNA damage. Stat: static, str: stretch, Vh: vehicle and inh: P38 inhibitor.

Since it had been reported that activation of P38-MAPK leads to an increase in P21, we inhibited P38-MAPK using a P38 inhibitor. However, when given the P38 inhibitor, there was a downregulation of P21 with stretch compared to the vehicle control (figure 6C), suggesting that P38 is involved in cellular senescence pathways in this *in vitro* model VILI. Nevertheless, inhibition of P38 did not decrease ɣH2AX protein level in the stretched SAEC (figure 6D), possibly due to DNA damage being upstream of P38-MAPK. It is also worth noting that there was decreased cell proliferation in all stretch groups, including stretch plus P38 inhibitor; this could be due to stretch causing DNA damage and regulating cell cycle via P21.

## Discussion

With the advent of novel respiratory diseases, including the current pandemic caused by COVID-19, more and more patients are ending up on mechanical ventilators, which lead to further injuries such as VILI. Despite being a lifeline for some patients, mechanical ventilators can leave lifelong damages [16] that are still not fully understood. In this study, we have developed both in vitro and in vivo models to simulate different damages caused by mechanical ventilation. We used these models to study the different mechanisms potentially involved in VILI, which can better understand the disease for future therapies.

### Damage due to MV

It has been extensively documented that the shear forces generated by mechanical ventilation exacerbate lung injury [17]. The characteristics of these lung injuries include the changes in lung structure. For instance, other studies, including ours, have shown that after mechanical ventilation, the airspace of mice becomes enlarged [18]–[20]. This is in line with what we observed in the lung histology of the mechanically ventilated mice. This airspace enlargement is also more pronounced in old mice generally, which is then exacerbated with mechanical ventilation (figure 1 A) due to the change in the lung’s parenchyma and alveolarization with age [21]. These structural changes in the lung could also lead to a change in the lung’s mechanics.

The structural damage and increased alveolar size, factors related to airspace enlargement, increase lung volume and compliance [22]. The increased alveolar space and the structural damage observed in their histology (figure 2) could explain why the old mice have inherently a higher lung single compartment static compliance compared to the young mice. However, other studies have shown that mechanical ventilation leads to a decrease in compliance [23], as observed in the young group in this study. Multiple studies have shown that structural lung damage leads to fluid exudation in the alveolar space [24]–[27]. This exudative phase is the cause of acute inflammation and leads to the release of the pro-inflammatory cytokine, which will then recruit immune cells to the site of the injury [28]. Amongst the recruited immune cells, PMN appears always to be recruited at a higher amount with ventilation. Bobba et al. have shown that PMNs are mechanosensitive, and they release cytokines in response to barotrauma, increased pressure usually associated with mechanical ventilation that causes alveolar damage [29]. The presence of PMN with mechanical ventilation correlate to our data where there was a high rise of PMN with ventilation in both young and old group.

### DNA damage and P21 in VILI

In addition to causing structural lung damage and recruitment of PMN, Blazquez-Prieto et al. have shown that when mechanically ventilated, mice exhibited a change in the nuclear envelope which coexisted with an increase ɣH2AX, a marker of DNA damage [7]. Moreover, increased accumulation of ɣH2AX has been associated with apoptosis [30] and cellular senescence [31]. Indeed, when cells sense DNA damage, they try to repair it by activating DNA damage response and increasing ɣH2AX, which could lead cells to cell cycle arrest and adapting senescence and a pro-inflammatory phenotype (SASP) [32] as a protective mechanism acutely to prevent further injury. However, activation of DNA damage response can also lead to apoptosis to clear out damaged cells [33]. Together with our results, where there was an increase in DNA damage manifested by increased ɣH2AX in both *in vitro* and *in vivo* experiments, these findings suggest that mechanical ventilation leads to early signs of senescence.

Strunz et al. have shown that epithelial cells in the alveolar are amongst the cells that experience DNA damage and apoptosis due to ventilation [14]. Apoptosis was further confirmed by TUNEL staining (figure 3), where mechanically ventilated mice (young and old) had larger total TUNEL positive cells than those that were not ventilated. The alveolar epithelium comprises two main cell types: AT1 and AT2 cells. During homeostasis and injury repair, AT2 cells are crucial to replacing AT1 cells which are terminally differentiated [13]. However, AT2 can also adopt a transient phenotype (which stain positive for KRT8) more susceptible to DNA damage. In this study, with mechanical ventilation and age, we observed an increase of KRT8 positive cells; but it was not until stretched that we had increased DNA damage with age. The susceptibility of these transient cells to DNA damage is evidence that mechanical ventilation can cause DNA damage, leading to a senescence-like phenotype.

P21 has been extensively linked to senescence and anti-proliferation [34]–[37]. In our model, the increased P21 gene expression and SASP manifested by increased IL6 with ventilation in both age groups indicate that mechanical ventilation here could be causing early senescence. In fact, during early senescence, the P21 level picks and declines as the cells are more committed to the senescence fate [38]. Another characteristic of senescent cells is a lack of proliferation due to cell cycle arrest. With increased senescence markers, there is a decrease in Ki67 [39]. This is in line with our findings where mechanical ventilation combined with age and age alone; there was a decrease in proliferation marked by a decrease in Ki67 (figure 4C); further signs that mechanical ventilation and age lead to senescence-like phenotype.

### The potential role for P38

TGF-β, which is involved in multiple physiological functions and diseases [40]–[43], has been shown to play an important role in pulmonary senescence [44], [45]. The sequestered TGF-β on the extracellular matrix is released during a stretch. Due to that release compounded with the injurious state of the environment that already exists, TGF-β becomes activated [46]. The activated TGF-β will cause a cascade of events, including activating the P38-MAPK pathway [47]. Activation of the P38-MAPK pathway plays an essential role in regulating the cell cycle by increasing P53, increasing P21 [48]. Though some parts of this mechanism have been extensible studied, the role of p38 inhibition has not been examined in an aging model of VILI. In the quest to investigate the mechanism of these senescence-like phenotypes in VILI, we used a P38 inhibitor to control the level of P21 observed with mechanical stretch ventilation. Inhibiting P38 in this study led to a decrease in P21 but not ɣH2AX because cyclic stretch leads to DNA damage that activates the P38-MAPK pathway.

### Limitations

The markers of senescence are very heterogeneous dynamic. Though markers traditionally associated with senescence were upregulated, we cannot be certain this is a chronic or irreversible phenotype as these markers can be found at other cellular states and conditions [49]. The hallmarks of senescence observed in this study could be a protective mechanism started by the cells to prevent further injuries [7]. They may revert when the acute injury is under control if the immune system has not already cleared out those cells. Further studies need to be done to confirm the state of senescence of the cells when stretched, such as survival study with long-term recovery. Though the study of the P38 mechanism performed on primary human lung cells isolated from the distal part of the alveoli provided some valuable information on how P21 is upregulated, more in vivo studies need to be performed t. We could not obtain cells isolated from young patients; this mechanism needs to be verified using cells from young patients to confirm if the mechanism holds with age.

## Conclusion

In conclusion, we have shown that high-pressure mechanical ventilation and age lead to structural damage at tissue, cellular, and proteins levels. These damages were correlated with the increase of KRT8 positive cells, which are susceptible to DNA damage and could play a vital role in senescence. We have also shown signs of senescence-like phenotypes with mechanical ventilation and with age. Finally, we have provided evidence that p38 may be a therapeutic target in stretch-induced senescence. These findings provide a better understanding of injuries resulting during mechanical ventilation and VILI and could be used for better-targeted therapies from injuries created by ventilation.

## Notes

### Competing Interest Statement

The authors have declared no competing interest.

